# Mammalian ANP32A and ANP32B proteins drive alternative avian influenza virus polymerase adaptations

**DOI:** 10.1101/2020.09.03.282384

**Authors:** Thomas. P. Peacock, Carol M. Sheppard, Ecco Staller, Rebecca Frise, Olivia C. Swann, Daniel H. Goldhill, Jason S. Long, Wendy S. Barclay

## Abstract

ANP32 proteins, which act as influenza polymerase co-factors, vary between birds and mammals. The well-known mammalian adaptation, PB2-E627K, enables influenza polymerase to use mammalian ANP32 proteins. However, some mammalian-adapted influenza viruses do not harbour this adaptation. Here, we show that alternative PB2 adaptations, Q591R and D701N also allow influenza polymerase to use mammalian ANP32 proteins. PB2-E627K strongly favours use of mammalian ANP32B proteins, whereas D701N shows no such bias. Accordingly, PB2-E627K adaptation emerges in species with strong pro-viral ANP32B proteins, such as humans and mice, while D701N is more commonly seen in isolates from swine, dogs and horses where ANP32A proteins are more strongly pro-viral. In an experimental evolution approach, passage of avian viruses in human cells drives acquisition of PB2-E627K, but not when ANP32B is ablated. The strong pro-viral support of ANP32B for PB2-E627K maps to the LCAR region of ANP32B.

## Introduction

The natural host reservoir of influenza A viruses is wild aquatic birds. To efficiently replicate in mammalian hosts, avian influenza viruses need to overcome multiple barriers by adapting to several mammalian host factors (Long, Mistry et al., 2019b). One major block to avian influenza virus replication in mammalian cells is the incompatibility of the viral polymerase with host acidic nuclear phosphoproteins of 32 kilodaltons (ANP32) proteins (Long, Giotis et al., 2016). ANP32 proteins are essential for influenza polymerase activity, and adaptation to the ANP32 proteins of a new host is generally the first mutation seen during cross-species jumps (Long et al., 2019b, Staller, Sheppard et al., 2019, Sugiyama, Kawaguchi et al., 2015, Zhang, Zhang et al., 2019). Most avian species encode ANP32A proteins that are longer than the mammalian orthologues due to an exon duplication that results in a 33 amino acid insertion between the N-terminal leucine rich repeat (LRR) domain and the C-terminal low complexity acidic region (LCAR). Avian influenza virus polymerase is supported by this longer avian-specific isoform of ANP32A, but not the shorter mammalian form (Long et al., 2016).

Mutations in the heterotrimeric viral polymerase can enable efficient use of mammalian ANP32 proteins, the best characterised of which is PB2-E627K (Domingues, Eletto et al., 2019, Long et al., 2016, Subbarao, London et al., 1993). However, some non-human mammalian influenza viruses do not contain PB2-E627K and have achieved mammalian adaptation through different mutations, for example the swine-origin H1N1 2009 pandemic virus (pH1N1) has PB2 polymorphisms at positions 271, 590 and 591 which functionally compensate for the lack of E627K (Liu, Qiao et al., 2012, Mehle & Doudna, 2009). Furthermore, all equine influenza virus strains, Eurasian avian-like swine influenza viruses, and canine influenza viruses lack PB2-E627K, but contain an alternative adaptation, PB2-D701N, previously described as modulating mammalian importin binding (Gabriel, Klingel et al., 2011, Sediri, Schwalm et al., 2015).

As well as differences in avian and mammalian ANP32 length, there also exists differences in the level of redundancy to support influenza virus polymerase in different vertebrate hosts. ANP32A is the sole ANP32 family member that supports influenza polymerase in birds, while in humans and most other mammalian influenza hosts ANP32A and ANP32B can both support polymerase activity to varying levels (Long, Idoko-Akoh et al., 2019a, Peacock, Swann et al., 2020, Staller et al., 2019, Zhang et al., 2019). One exception is mice in which only ANP32B can efficiently support influenza polymerase due to a polymorphism in murine ANP32A at position 130 – a residue critical for the interaction between ANP32 proteins and viral polymerase (Beck, Zickler et al., 2020, Staller et al., 2019, Zhang et al., 2019). It has recently been shown that, although both swine ANP32A and ANP32B can support mammalian-adapted polymerases, swine ANP32A has the more potent pro-viral function and can even partially support avian polymerases (Peacock et al., 2020, Zhang, Li et al., 2020). Similarly in horses, dogs, seals and bats, ANP32A appears to be a more potent pro-viral factor than ANP32B (Peacock et al., 2020).

In this study we aimed to understand whether the alternative mammalian PB2 adaptations, other than PB2-E627K, function by adapting the polymerase to utilise mammalian ANP32 proteins, or whether they achieve adaptation through an ANP32-independent mechanism. We find that alongside PB2-E627K, -Q591R and -D701N specifically adapt avian polymerases to use mammalian ANP32 proteins. Furthermore, while PB2-D701N allows the polymerase to efficiently use both ANP32A and B proteins, -E627K specifically favours use of mammalian ANP32B proteins. In support of this finding we use bioinformatics to show that viruses adapting to hosts in which ANP32B is the more potent pro-viral factor, such as humans and mice, generally adapt via PB2-E627K while viruses that emerge in pigs, horses or dogs, hosts with potent pro-viral ANP32A proteins, more often gain PB2-D701N, or -Q591R. Using an experimental evolution approach we passaged a pair of avian influenza virus in human cells and observed the emergence of the PB2-E627K mutation. This adaptation was not seen during passage of the same avian virus in human cells lacking ANP32B (BKO), or in swine cells with a dominant ANP32A protein. Finally, we map the difference in ANP32B preference of polymerases containing E627K to the ANP32B protein LCAR, the region that has recently been implied to directly interact with the 627 domain in the influenza polymerase/ANP32 co-structure (Camacho-Zarco, Kalayil et al., 2020).

## Results

### PB2-Q591R, -E627K and -D701N specifically adapt avian influenza polymerases to mammalian ANP32 proteins

We previously described a library of mammalian PB2 adaptations in an avian influenza virus polymerase backbone A/turkey/England/50-92/1991(H5N1; 50-92) (Cauldwell, Moncorge et al., 2013). Here we expanded the library to include some additional mutants implicated in the literature as mammalian-adapting variants (Table 1). We tested the effect of each mutation on polymerase activity in wild-type (WT) human eHAP and chicken DF-1 cells (Figure 1). Consistent with our previous findings (Cauldwell et al., 2013), mutations displayed one of three phenotypes: i) no significant increase in human or avian cells (G590S -grey bars), ii) significantly increased polymerase activity in both human and avian cells (G158E, T271A, K702R and D740N – red bars), or iii) significantly increased polymerase activity only in human cells (Q591R, E627K/V and D701N – green bars). This implies that the third set of PB2 mutants work by adapting the polymerase to utilising a host factor that is different between mammalian and avian cells.

**Table 1.** Summary table of PB2 mutants used in this study and effect shown by mutants with ANP32 proteins

| Mutation | Group | Impact |
| --- | --- | --- |
| G158E | Group 2 | Non species-specific polymerase activity boost |
| T271A | Group 2 | Non species-specific polymerase activity boost |
| G590S | Group 1 | Little or no effect alone |
| Q591R | Group 3 | Mammalian ANP32-specific adaptation (non-biased) |
| E627K | Group 3 | Mammalian ANP32-specific adaptation (ANP32B-biased) |
| E627V | Group 3 | Mammalian ANP32-specific adaptation (ANP32B-biased) |
| D701N | Group 3 | Mammalian ANP32-specific adaptation (non-biased) |
| K702R | Group 2 | Non species-specific polymerase activity boost |
| D740N | Group 2 | Non species-specific polymerase activity boost |

**Figure 1.**
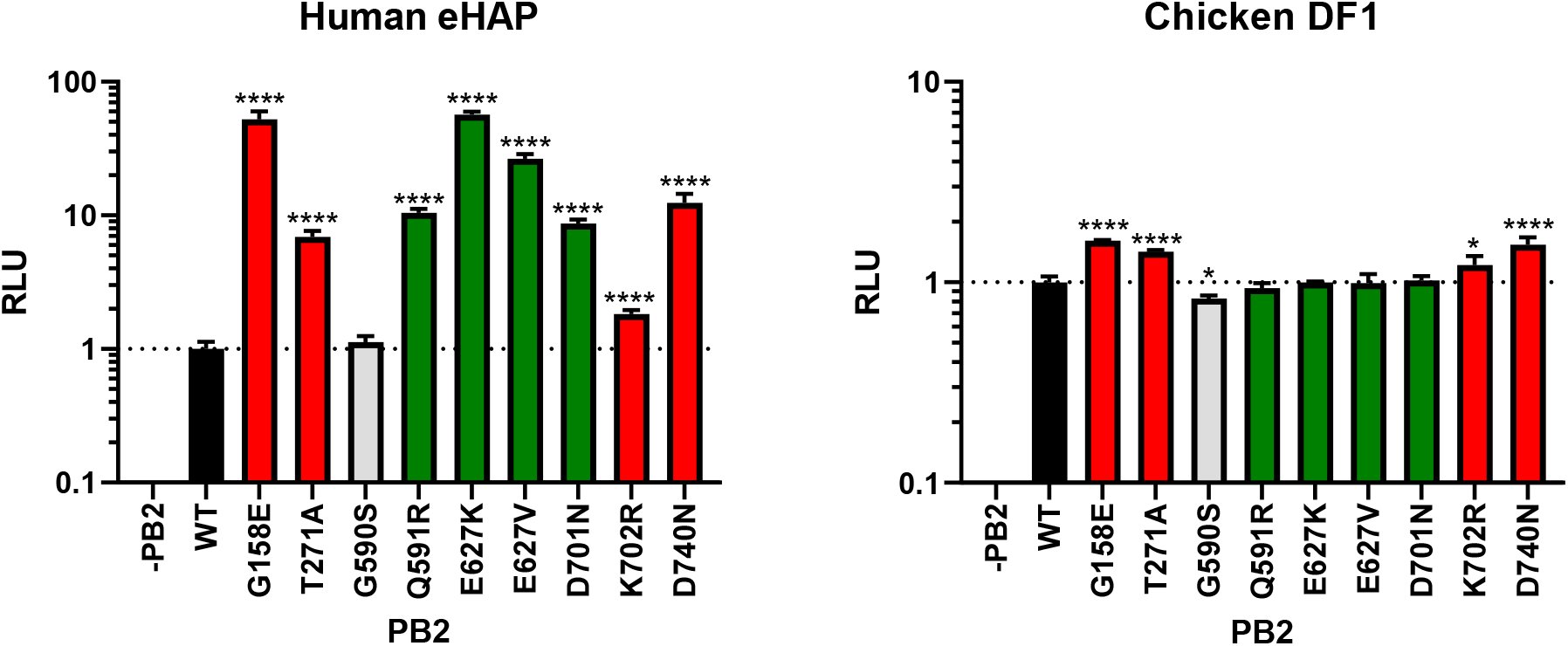
PB2-G158E, T271A, K702R and D740N increase polymerase activity in a non-mammalian specific manner. Minigenome assays performed in WT human eHAP cells or WT chicken DF-1 cells with avian 50-92 polymerase with different mammalian adaptations. Data throughout indicates triplicate repeats plotted as the mean and standard deviation, normalised to PB2 WT. Statistical significance was determined by one-way ANOVA with multiple comparisons against WT on log-transformed data. Lognormality of data was confirmed by Shapiro-Wilk test of normality. *, 0.05 ≥ P > 0.01; **, 0.01 ≥ P > 0.001; ***, 0.001 ≥ P > 0.0001; ****, P ≤ 0.0001.

A range of host factors have been implicated in the mammalian adaptation of avian influenza virus polymerase including α-importins, DDX17 and ANP32 proteins (reviewed in (Long et al., 2019b)). To investigate whether the PB2 mutations in our panel adapted polymerase to mammalian ANP32 proteins, we performed an ANP32 complementation assay in human cells lacking endogenous ANP32A and ANP32B (dKO) (Long et al., 2019a, Staller et al., 2019). We tested the complementation of polymerase activity for each PB2 mutant with ANP32 proteins from chicken, human, swine, or dog (Figure 2A, Supplementary Figure S1).

**Figure 2.**
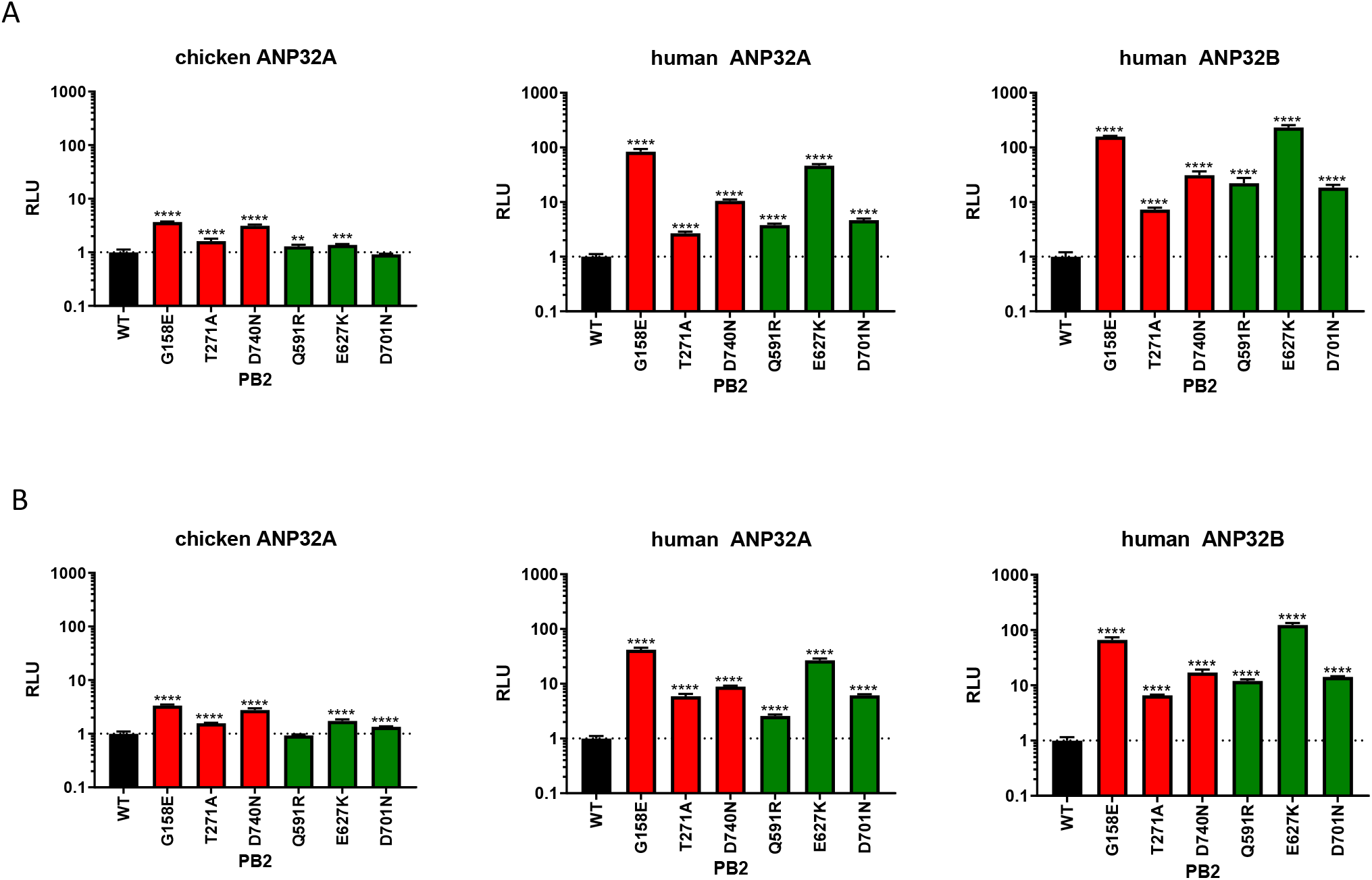
PB2-Q591R, -E627K and -D701N specifically adapt influenza virus polymerase to human ANP32 proteins. Minigenome assays performed in (A) human eHAP dKO cells or (B) chicken DF-1 AKO cells with avian 50-92 polymerase with different mammalian adaptations transfected in along with different chicken or human ANP32 proteins. Data throughout indicates triplicate repeats plotted as the mean and standard deviation, normalised to PB2 WT. Statistical significance was determined by one-way ANOVA with multiple comparisons again WT on log-transformed data. Lognormality of data was confirmed by Shapiro-Wilk test of normality. *, 0.05 ≥ P > 0.01; **, 0.01 ≥ P > 0.001; ***, 0.001 ≥ P > 0.0001; ****, P ≤ 0.0001.

As before the PB2 mutants displayed one of three phenotypes that corresponded with their phenotype in human or chicken cells described in Figure 1. Mutant group 1 were rescued by co-expression of chANP32A but not by expression of mammalian ANP32A or B, and gave patterns of polymerase activity largely identical to WT PB2 (Supplementary Figure S1A – grey bars), activity of group 2 mutants was significantly better than WT PB2 by co-expression of either chicken or mammalian ANP32 (Figure 2A – red bars), and group 3 mutants were only enhanced over WT in presence of mammalian ANP32 proteins (Figure 2A – green bars; Supplementary Figure S1A). This implies that the PB2 mutations at amino acids 591, 627 and 701 all enhance polymerase activity in mammalian cells by specifically enabling complementation by mammalian ANP32 proteins.

We then considered whether any other host specific factors that differed between avian and mammalian cells might affect the complementation of influenza polymerase by ANP32 proteins. We reconstituted each polymerase containing different PB2 mutations in chicken DF-1 cells in which chANP32A expression was ablated by CRISPR editing (Long et al., 2019a), and then rescued polymerase activity by again co-expressing ANP32 proteins from chicken, human, swine or dog (Figure 2B, Supplementary Figure S1B). Overall, the pattern of complementation in chicken cells was consistent with that seen in human cells, suggesting that mammalian adapting polymerase mutations in PB2 enable mammalian ANP32 proteins to support polymerase activity even in chicken cells, and no other host factors that differ between avian and mammalian species are required for this phenotype. In either human and chicken cells lacking ANP32 proteins and complemented with chicken ANP32A, group 3 PB2 mutations Q591R, E627K, and D701N did give a small boost to polymerase activity, however this boost was far less than that seen for the group 2 mutants (Figure 2, Supplementary Figure 1).

These results indicate that PB2 mutations Q591R and D701N, but not the other mutations tested here, act in a similar manner to E627K and enable viral polymerase to utilise mammalian ANP32 proteins as a co-factor.

### PB2-E627K, but not -D701N, adapts influenza virus polymerase for preferential complementation by mammalian ANP32B

Across different mammalian species, there is variation in the ability of ANP32A or ANP32B to support influenza virus polymerase. For example, in humans and mice ANP32B is the more potent polymerase co-factor, whilst the ANP32A protein of pigs, horses, dogs, seals and bats more efficiently support polymerase activity (Supplementary Figure S2A,B)(Peacock et al., 2020, Zhang et al., 2019). This dominance is maintained across a range of different ratios of these proteins (Supplementary Figure S2C). These subtle variations are due to polymorphisms between the ANP32 orthologues in different species. For example, the potent pro-viral activity of swine ANP32A is due to polymorphisms at positions 106 and 156 (Peacock et al., 2020, Staller et al., 2019, Zhang et al., 2020). Similarly, the weak pro-viral activity of dog (as well as bat and seal) ANP32B is attributed to residue 153 that is glutamine in the human ANP32B, but arginine in the canine orthologue (Supplementary Figure S3A-C)

To further investigate the compatibility between different polymerase constellations and different mammalian ANP proteins, polymerase reconstituted with PB2 mutants from group 3 were tested to see if they displayed any bias in ANP32 paralogue usage (Figure 3). The relative efficacy of each ANP32 protein to support polymerase varied depending on the nature of adaptive mutation in PB2. For polymerase bearing PB2-D701N, human ANP32B was superior to human ANP32A, whereas swine ANP32A was more supportive than ANP32B and canine ANP32B was poorly supportive. In contrast, for polymerase bearing E627K both human and swine ANP32B were more potent than their respective ANP32A counterparts. Moreover, E627K polymerase was also somewhat supported by canine ANP32B, which is very poorly used by most polymerases (Supplementary Figure S2A)(Peacock et al., 2020). PB2-E627K was more highly supported by human ANP32B than ANP32A over a wide range of plasmid concentrations (Supplementary Figure 4). For polymerase bearing PB2-Q591R the pattern was intermediate: little difference was seen in swine ANP32 preference while human ANP32B was significantly preferred over ANP32A and dog ANP32B was ineffective as a proviral factor. These effects were observable in ANP32 complementation assays in both human and chicken cells (Figure 3, Supplementary Figure S5).

**Figure 3.**
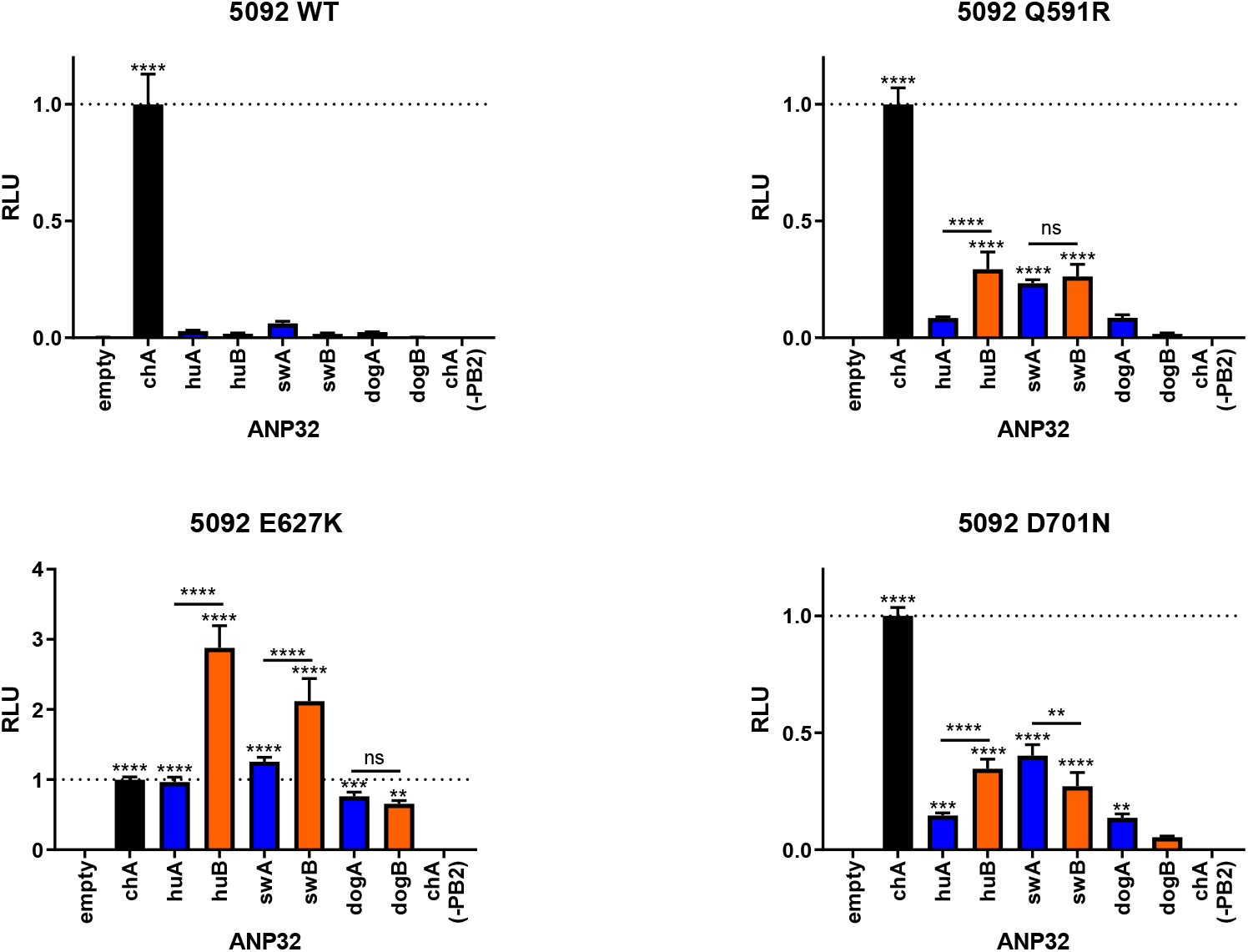
PB2-E627K shows a greater preference than -Q591R or -D701N for using mammalian ANP32B proteins. Data from Figure 1 and Supplementary Figure S1 shown in a different format. Minigenome assays performed in human eHAP dKO cells with avian 50-92 polymerase with different mammalian adaptations transfected in along with different avian or mammalian ANP32A (blue bars) or ANP32B proteins (orange bars). Data throughout indicates triplicate repeats plotted as the mean and standard deviation, normalised to chicken ANP32A. Statistical significance was determined by one-way ANOVA with multiple comparisons, statistical tests without comparison bars indicate a comparison against empty vector and between ANP32A and ANP32B proteins from the same species. *, 0.05 ≥ P > 0.01; **, 0.01 ≥ P > 0.001; ***, 0.001 ≥ P > 0.0001; ****, P ≤ 0.0001.

**Figure 4.**
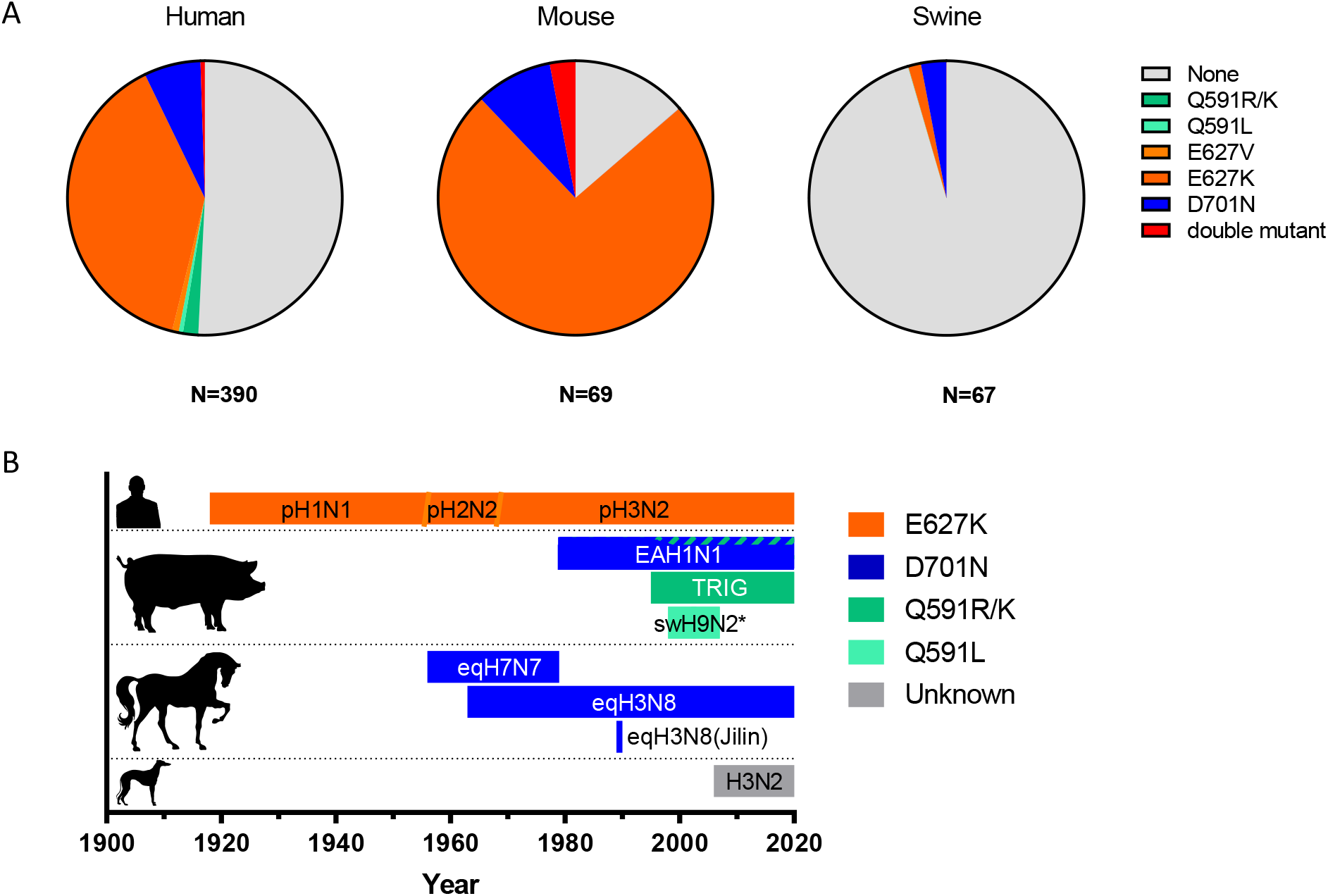
Mammalian species with dominantly pro-viral ANP32B, but not ANP32A proteins, are associated with E627K. Mammalian adaptations seen in avian-origin viruses during (A) zoonotic or likely dead-end cross-species infections/mouse passage experiments (middle panel only) and (B) stable circulation and prolonged adaption to a mammalian host. Human and swine cross-species infections (A, left and right panel) were calculated by downloading all non-H1-3 human/swine influenza virus strain PB2s from NCBI, performing an alignment, curating out any seasonal human influenza segments and looking at the identities of position 591, 627 and 701. Mouse adaptation studies (A, middle panel) were calculated by performing a literature review of any influenza mouse adaptation studies using avian-origin influenza virus without any prior mammalian adaptation. Only virus strains with good evidence for being stably circulating in their respective mammalian species were included in the timeline (B). *swine H9N2 was included as the phylogenetic and molecular evidence was strong that, although this virus does appear to potentially co-circulate in both swine and chickens, the virus does appear to show some clear mammalian adaptation markers.

These data illustrate a bias of different PB2 polymerase adaptations for different mammalian ANP32A or B proteins that is particularly evident for PB2-E627K, and to a lesser extent, -Q591R, which show a preference for ANP32B proteins.

### Species with strongly proviral ANP32B proteins drive the acquisition of PB2-E627K

Although PB2-E627K is the key polymerase adaptation in human seasonal influenza virus and commonly found in human zoonotic infections as well as in laboratory-adapted mouse-passaged avian influenza viruses, it is rarely found in viruses that have crossed from birds into swine, dogs or horses (Liu et al., 2012, Lloren, Lee et al., 2017). Instead, viruses endemic in those species tend to harbour the other ANP32-specific mammalian adaptations, PB2-Q591R and -D701N. We therefore hypothesised that in humans, PB2-E627K might evolve as a specific adaptation to the strongly proviral ANP32B. Conversely for other mammalian species such as dogs and horses, the selective pressure exerted by ANP32B would be weaker and adaptation would likely occur by PB2-D701N.

To test this hypothesis, we first performed bioinformatics analysis comparing the number of mammalian adaptions found at sites 591, 627 and 701 during zoonotic infections, laboratory mouse adaptation studies, or sustained transmission of avian-origin PB2 segments in different mammalian species (Figure 4A). Avian influenza viruses strongly selected for PB2-E627K during both human zoonotic events and mouse experimental adaptation, although other ANP32-specific mutations were also represented, such as PB2-D701N. Incursions of avian influenza viruses into pigs rarely showed adaptation at any of these three sites (Figure 4A, right panel). This might be explained by the observation that swine ANP32A is somewhat supportive of non-adapted avian influenza virus polymerases (Peacock et al., 2020, Zhang et al., 2020). It is also noteworthy that the number of viruses with multiple ANP32-specific PB2 adaptations simultaneously is very low, suggesting these mutations are partially redundant (although the more poorly adaptive Q591R/K and D701N mutations were occasionally found together).

In viruses that have crossed from birds and sustainably circulated in mammalian hosts we saw a clear difference in PB2 adaptations in humans, compared to viruses endemic in swine, horses and dogs. The only sustained avian-origin PB2 in humans (which transmitted during the 1918 Spanish influenza pandemic) has E627K, whereas none of the polymerases from viruses adapted in other species show this adaptation. Instead such viruses contain a mixture of D701N and Q591R/K/L or (in the case of canine H3N2 viruses) none of the currently described mammalian ANP32-specific adaptations (Figure 4B).

### Experimental evolution of an avian influenza virus in human cells shows expression of ANP32B leads to PB2-E627K

To investigate whether different ANP32 proteins drive different PB2 adapting mutations, we used an experimental evolution approach, serially passaging an avian influenza virus though human cell lines lacking either ANP32A (AKO) or ANP32B (BKO) (Staller et al., 2019).

Six populations of avian influenza virus 50-92 were passaged 10 times through either control cells (which express both ANP32A and ANP32B), or cells that express ANP32B only (AKO) or ANP32A only (BKO). PB2 segments from each population were Sanger sequenced at passages 2, 5 and 10. By passage 5, in both the control and AKO cells, three out of six (50%) of the populations had evolved PB2-E627K. This adaptation was not detected in the BKO cells, even by passage 10. Instead in these cells one virus population (17%) at passage 5, and two by passage 10 (33%), gained the D701N mutation (Figure 5A). D701N was also seen in one or two populations in WT or AKO cells by passage 10, respectively.

**Figure 5.**
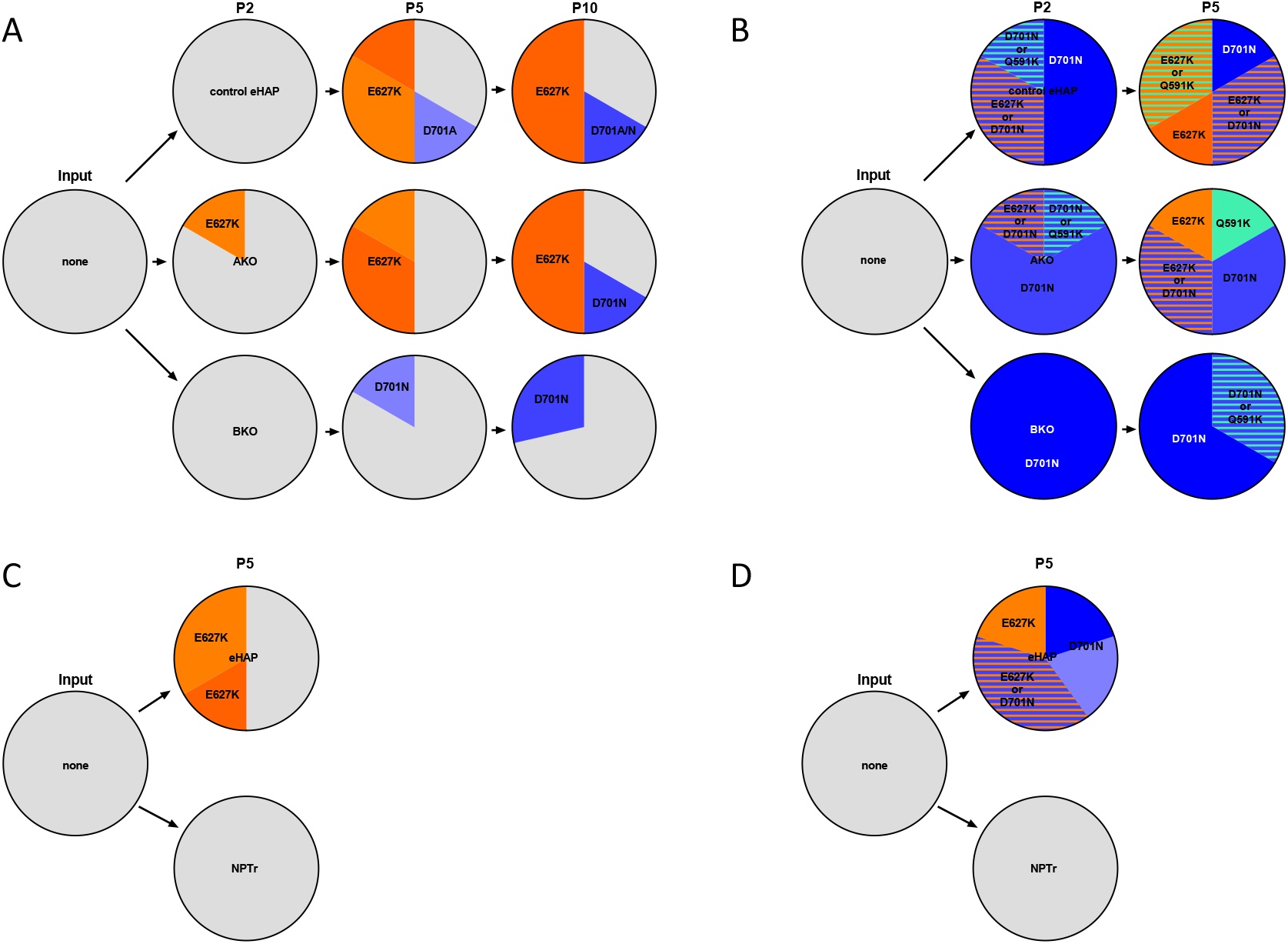
Experimental evolution of an avian influenza virus in human cells abrogated for ANP32B does not lead to the PB2-E627K adaptation. Sequencing summary of avian origin 50-92 (A, C) or Anhui (B, D) in human cells (A, B) ablated for ANP32A or ANP32B or (C, D) WT human and swine cells. Each pie chart indicates 6 independently passaged populations, depth of colour indicates rough estimation of the proportion of the population – light shades indicate a mixed population of WT and indicated residues at the position while darker shades indicate a complete change. Striped bars indicate mix of both E627K and D701N. Grey slices indicate no changes are detectible at positions 591, 627 or 701.

To confirm this phenotype was not specific to a single strain of avian influenza, this experiment was repeated with an H7N9 virus, A/Anhui/1/2013 (Anhui). The Anhui isolate had naturally gained PB2-E627K during zoonosis but this was reverted to 627E by reverse genetics for the purpose of the passage experiment. In a similar manner to the H5N1 50-92 virus, the Anhui PB2 gene gained a mixture of E627K, D701N, as well as Q591K, in the control cells or AKO cells by passage 5, whereas in the BKO cells every population evolved D701N or Q591K but E627K was not detected (Figure 5B).

The same pair of avian influenza viruses were also passaged in swine origin NPTr cells. In stark contrast to the human cells, no ANP32 adaptations were seen after 5 passages in the swine NPTr cells (Figure 5C,D). Taken together, these observations suggest that the predominance for the PB2-E627K adaptive mutation seen in human cells is driven by adaptation to utilise human ANP32B.

### The PB2-E627K preference for human ANP32B maps to the LCAR domain of the protein

To investigate the molecular basis of the superior ability of human ANP32B over ANP32A to complement mammalian-adapted PB2-E627K polymerase, we generated human ANP32A/B chimeric ANP32 constructs. As the LCAR is described as directly interacting with the 627 domain of PB2 (Camacho-Zarco et al., 2020, Mistry, Long et al., 2019), we switched the LCAR between human ANP32A and B (from amino acid 161 to the C-terminus, red highlight - Figure 6A). Human ANP32A with the ANP32B LCAR was much more efficient at rescuing PB2-E627K polymerase activity than WT human ANP32A, while conversely introducing the ANP32A LCAR onto human ANP32B reduced its capacity to support polymerase activity despite robust expression of the chimeric proteins (Figure 6B,C). This pattern held true for a pair of unrelated H1N1 and H5N1 avian-origin polymerases tested (Figure 6B). This suggests that the preference for ANP32B shown by PB2-E627K polymerase maps to amino acid differences in the ANP32 LCAR domain and provides further evidence that this ANP32 domain likely directly interacts with the PB2 627/NLS domain.

**Figure 6.**
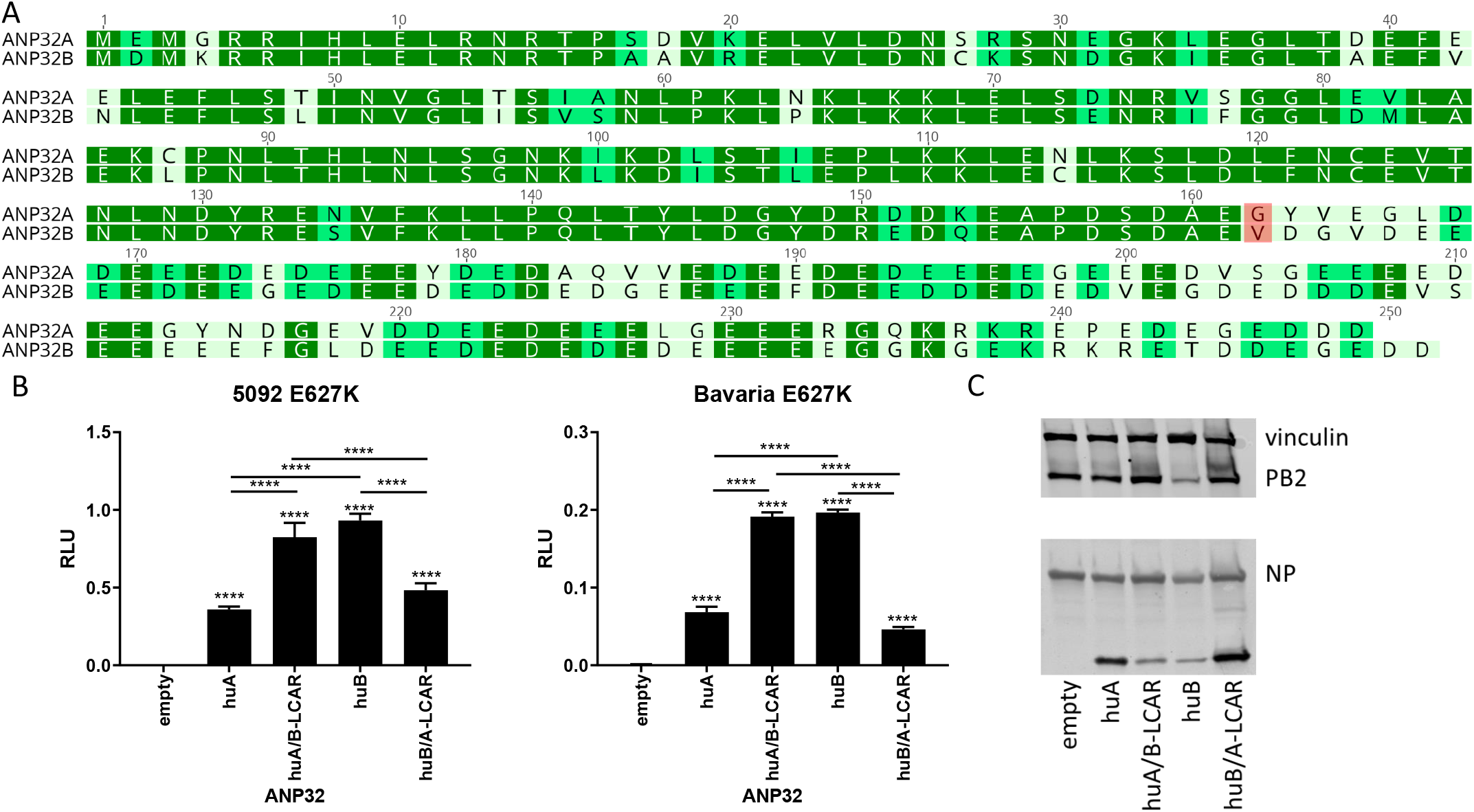
Differences in the LCAR of human ANP32A and ANP32B are responsible for the preference of PB2-E627K viruses for ANP32B. (A) Non-aligned comparison of human ANP32A and ANP32B sequences indicating sites of interest in yellow and blue. (B) Minigenome assays performed in human eHAP dKO cells with avian 50-92 or Bavaria polymerase with different mammalian adaptations transfected in along with different chimeric human ANP32 protein. Data throughout indicates triplicate repeats plotted as the mean and standard deviation. Statistical significance was determined by one-way ANOVA with multiple comparisons, statistical tests without comparison bars indicate a comparison against empty vector and between the different ANP32 proteins. ****, P ≤ 0.0001. (C) Western blot of chimeric human ANP32 proteins used in part (B).

## Discussion

In this study we showed that several different mutations in PB2 avian-origin influenza polymerases to use the shorter ANP32 proteins found in mammalian cells but that the most well-known of these PB2-E627K, strongly biases polymerases towards reliance on mammalian ANP32B. While ANP32A and ANP32B serve redundant proviral roles in many mammals, ANP32B is the dominant proviral factor in humans and mice whereas in most other relevant mammalian hosts, such as pigs, horses and dogs, ANP32A proteins is the more potent (Peacock et al., 2020, Staller et al., 2019, Zhang et al., 2020, Zhang et al., 2019). This pattern shapes the adaptive evolution of avian viruses in these different mammalian hosts. Thus, adaptation in humans and mice tends to strongly select for PB2-E627K while pigs, dogs and horses tend to select for PB2-D701N, or Q591R/K. This was borne out in an experimental evolution study where WT human cells, or those lacking ANP32A drove viruses to gain PB2-E627K, whereas viruses passaged through cells lacking ANP32B did not acquire that mutation. Finally we find that the strong preference for ANP32B proteins granted by PB2-E627K is due to differences between ANP32A and ANP32B sequence in the LCAR region of these proteins, a region implicated with direct interaction with the PB2 627 and NLS domains (Camacho-Zarco et al., 2020, Mistry et al., 2019). Overall these data suggest that the evolutionary ecology of influenza virus polymerase differs in different species due to differences in mammalian ANP32 proteins (Figure 7).

**Figure 7.**
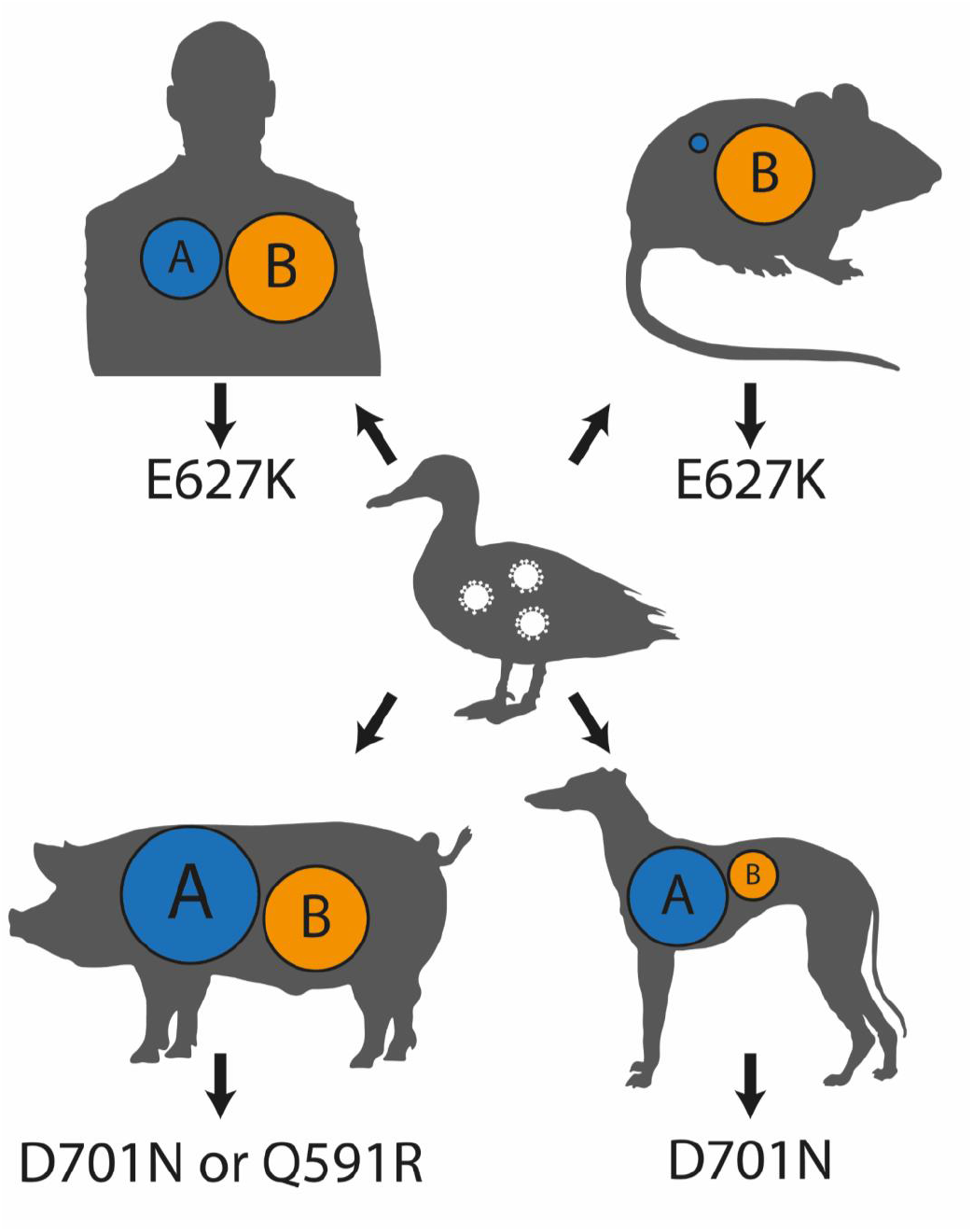
Model of how ANP32 protein dominance in different species may drive mammalian adaptation. Summary model of the data from this study showing how avian-origin influenza viruses adapt to the dominant pro-viral ANP32 protein in a new host and gain different adaptive mutation in PB2. Size of circles in each host indicate the relative pro-viral ability of those ANP32 proteins.

Previous studies have shown that varying expression of ANP32A isoforms in different avian species result in a different propensity to drive mammalian-like polymerase adaptations (e.g. PB2-E627K)(Baker, Ledwith et al., 2018, Domingues et al., 2019). Our work further expands on this concept that the pattern of ANP32 expression in a species can influence virus evolution. This finding may have implications for the relative risk of zoonotic or pandemic viruses emerging from different species; for example, although humans are often exposed to equine and canine hosts there is a lack of evidence for zoonotic influenza infections originating from these species, although mismatched HA receptor binding preference is also likely to contribute to this interspecies block (Collins, Vachieri et al., 2014).

Surprisingly, we found that PB2-D701N was an efficient adaptation to short mammalian ANP32 family members, regardless of whether this assay was undertaken in human or chicken cells. Previously PB2-D701N has been implicated as an adaptation to a different set of host factors, the importin-α family (Gabriel et al., 2011, Sediri et al., 2015). Our results from avian cells would imply that importin-α adaptation is not the only phenotype of this mutation, since, in the presence of chicken importins, polymerases reconstituted with this mutation were efficiently supported by mammalian ANP32 proteins.

Influenza virus polymerases co-opt a range of host factors for their replication and transcription (Peacock, Sheppard et al., 2019). Our approach of using human and avian cells lacking pro-viral ANP32 family members has potential as a powerful screening method for investigating, less well-defined mammalian ANP32 adaptations, and to discover novel host factors that affect polymerase activity including proviral and restriction factors in human cells (Mehle & Doudna, 2008). Future work should further map the sequences in the ANP32B LCAR that direct the evolution of the PB2-E627K mutations and attempt to understand the preference for different PB2 adaptations driven by ANP32A or B from a structural perspective.

## Materials and methods

### Cells

Human engineered-Haploid cells (eHAP; Horizon Discovery) without gene knockout (Control) or with ANP32A (AKO), ANP32B (BKO), or with both knocked out (dKO) by CRISPR-Cas9, as described previously (Staller et al., 2019), were maintained in Iscove’s Modified Dulbecco’s Medium (IMDM; ThermoFisher) supplemented with 10% faetal bovine serum (FBS; Biosera), 1% non-essential amino acids (NEAA; Gibco) and 1% Penicillin-streptomycin (pen-strep; invitrogen). Human embryonic kidney (293Ts, ATCC), Madin-Darby canine kidney cells (MDCK, ATCC) and swine Newborn Pig Trachea cells (NPTr; ATCC), were maintained in Dulbecco’s Modified Eagle Medium (DMEM) supplemented with 10% FBS, 1% NEAA and 1% pen-strep. Chicken fibroblast (DF-1; ATCC) were maintained in DMEM supplemented with 10% FBS, 5% tryptose phosphate broth (Sigma), 1% NEAA and 1% pen-strep. All mammalian cells were maintained at 37°C, 5% CO_2_ while DF1s were maintained at 39°C, 5% CO_2_.

### Plasmid constructs

Viruses and virus minigenome full strain names used through this study were A/duck/Bavaria/1/1977(H1N1, Bavaria), A/turkey/England/50-92/1992(H5N1; 50-92), A/Anhui/1/2013(H7N9; Anhui), A/England/687/2010(pH1N1), A/Victoria/1975(H3N2), and A/swine/England/453/2006(H1N1). Viral minigenome expression plasmids were either generated previously or made using overlap extension PCR (Cauldwell et al., 2013, Elderfield, Watson et al., 2014, Moncorge, Long et al., 2013). ANP32 expression constructs were made as previously described or generated using overlap extension PCR (Long et al., 2016, Peacock et al., 2020, Staller et al., 2019).

### Virus strains

All virus work in this study was performed with A/turkey/England/50-92/1992(H5N1; 50-92) or A/Anhui/1/2013(H7N9) [K627E], reassortant viruses was generated by rescuing the polymerase, NP and NS segments of the homologous virus with the HA, NA and M segments of A/Puerto Rico/1/1934(H1N1; PR8) as previously described (Long, Howard et al., 2013). Virus was titred by plaque assay on MDCKs. Virus PB2s were sequenced to confirm no prior mammalian adaptation was acquired during rescue or propagation. Infections were carried out at 37°C in relevant virus-containing serum-free media (DMEM or IMDM, 1% NEAA, 1% P/S). 1 hour after infection media was changed to serum-free media supplemented with 1µg/ml tosyl phenylalanyl chloromethyl ketone (TPCK)-treated trypsin (Worthington-Biochemical). Passage experiments were performed by infecting cells at an MOI of 0.01, 48 hours after inoculation viruses were harvested and prepared for the next passage.

### Minigenome assay

eHAP dKO cells were transfected at around 50% confluence in 24 well plates using lipofectamine® 3000 (thermo fisher) with the following mixture of plasmids; 100ng of pCAGGs ANP32 or empty pCAGGs, 40ng of pCAGGs PB2, 40ng of pCAGGs PB1, 20ng of pCAGGs PA, 80ng of pCAGGs NP, 40ng of pCAGGs Renilla luciferase, 40ng of polI vRNA-Firefly luciferase. DF-1 AKO cells were transfected in 12 well plates using double the amount of plasmid in eHAP cells and polI vRNA-firefly plasmids with a chicken polI site described previously (Moncorge, Mura et al., 2010). 24 hours post-transfection cells were lysed in passive lysis buffer (Promega) and luciferase bio-luminescent signals were read on a FLUOstar Omega plate reader (BMG Labtech) using the Dual-Luciferase® Reporter Assay System (Promega). Firefly signal was normalised to *Renilla* signal to give relative luminesce units (RLU). All assays were performed on a minimum of two separate occasions with representative data shown.

### Western Blotting

To visualise protein expression during mini-genome assays, around 500,000 transfected cells were lysed in RIPA buffer (150mM NaCl, 1% NP-40, 0.5% Sodium deoxycholate, 0.1% SDS, 50mM TRIS, pH 7.4) supplemented with an EDTA-free protease inhibitor cocktail tablet (Roche).

Proteins were detected with mouse α-FLAG (F1804, Sigma), rabbit α-Vinculin (AB129002, Abcam), rabbit α-PB2 (GTX125926, GeneTex) and mouse α-NP ([C43] ab128193, Abcam). The following near infrared (NIR) fluorescently tagged secondary antibodies were used: IRDye® 680RD Goat Anti-Rabbit (IgG) secondary antibody (Ab216777, Abcam) and IRDye® 800CW Goat Anti-Mouse (IgG) secondary antibody (Ab216772, Abcam). Western Blots were visualised using an Odyssey Imaging System (LI-COR Biosciences).

### Experimental virus evolution

At each passage 1000 pfu of avian influenza virus was inoculated in serum-free media into confluent monolayers seeded in 6 well plates. After 1 hour, media was replaced with serum-free media with 1ug/ml of TPCK trypsin. 48 hours post-infection the supernatant was harvested, spun down to remove cellular debris and used for further passages. Samples were sequenced (where appropriate), and frozen down and stored at -80°C. Each passage experiment was done with 6 concurrent populations.

### Virus sequencing

To sequence viruses, RNA was extracted from cell-free virus-containing supernatants using the viral RNA extraction mini kit (Qiagen). cDNA synthesis was conducted using Superscript IV and the uni12-FluG primer (AGCGAAAGCAGG). Sequencing of PB2s were performed using two sets of primers with 5’-M13F or M13R primer sites (TGTAAAACGACGGCCAGTCCACTGTGGACCATATGGCC with CAGGAAACAGCTATGACCTGGAATATTCATCCACTCCC, and TGTAAAACGACGGCCAGTGGGAGTGGATGAATATTCCAG with CAGGAAACAGCTATGACCGCTGTCTGGCTGTCAGTAAGTATGC). PA was sequenced similarly (TGTAAAACGACGGCCAGTGCGACAATGCTTCAATCCAATG with CAGGAAACAGCTATGACCCTTCTCATACTTGCAATGTGCTC, and TGTAAAACGACGGCCAGTGGGCACTCGGTGAGAACATGGC with CAGGAAACAGCTATGACAACTATTTCAGTGCATGTG). PCR was performed using KOD Hot Start DNA polymerase (Merck). PCR products were purified using the Monarch PCR and DNA Cleanup Kit (NEB) and sequenced using the Sanger method with M13F or M13R primers.

### Bioinformatics analysis and literature review

To assess the proportion zoonotic influenza viruses with different mammalian ANP32 adaptations the PB2 sequences of all non-H1, -H2 and -H3 human or swine influenza viruses were downloaded from the NCBI influenza virus database and aligned. Sequences that were determined to be of seasonal human influenza origin were identified by BLASTn and removed and proportions of viruses with adaptation at position 591, 627 and 701 were calculated. For the mouse adaptation summary, an exhaustive literature search was undertaken for any study taking an avian-origin virus without any prior mammalian adaptation and passaging it through mice in pubmed using the search terms “mouse”, “influenza” and either “adaptation”, “adaption” or “passage”. A list of the papers identified and included in this analysis is included in supplementary table S1.

For the timeline of stably circulating avian-origin mammalian influenza virus strains – viruses were chosen due to the strength of the evidence that they came directly from birds into said species and not from another mammalian species – hence why pH1N1 swine-origin H1N1 and equine-origin canine H3N8 are excluded. Swine H9N2 was selected to be included as there remains fairly good evidence that, although this virus may have continued to co-circulate between poultry and pigs, it does show several mammalian adaptations and therefore probably constituted a swine-adapted avian-origin virus.

### Safety and Biosafety

All studies of infectious agents were conducted within biosafety level 2 facilities approved by the UK Health and Safety Executive and in accordance with local rules at Imperial College London.

### Data availability

This manuscript contains no data deposited in external repositories.

## Acknowledgements and funding

The authors would like to thank members of the Barclay lab of Imperial College London for their scientific input, advice, and support for this project.

T.P.P. was supported by BBSRC grant BB/R013071/1; C.M.S, D.H.G. and W.S.B were supported by Wellcome Trust grant 205100; E.S. was supported by an Imperial College President’s Scholarship; R.F. was supported by Wellcome Trust grant 200187; O.C.S. was supported by a Wellcome Trust studentship. J.S.L. and W.S.B were supported by BBSRC grant BB/K002465/1; W.S.B was supported by BBSRC grant BB/S008292/1.

## Author Contributions

TPP and WSB conceived this study. TPP, CMS, ES, RF, OCS performed the experiments. TPP and DHG performed data curation and analysis. JSL and WSB acquired funding for the study. TPP and WSB wrote the manuscript. TPP, CMS, ES, OCS, DHG, JSL and WSB reviewed and edited the paper.

---

**Figure S1.**
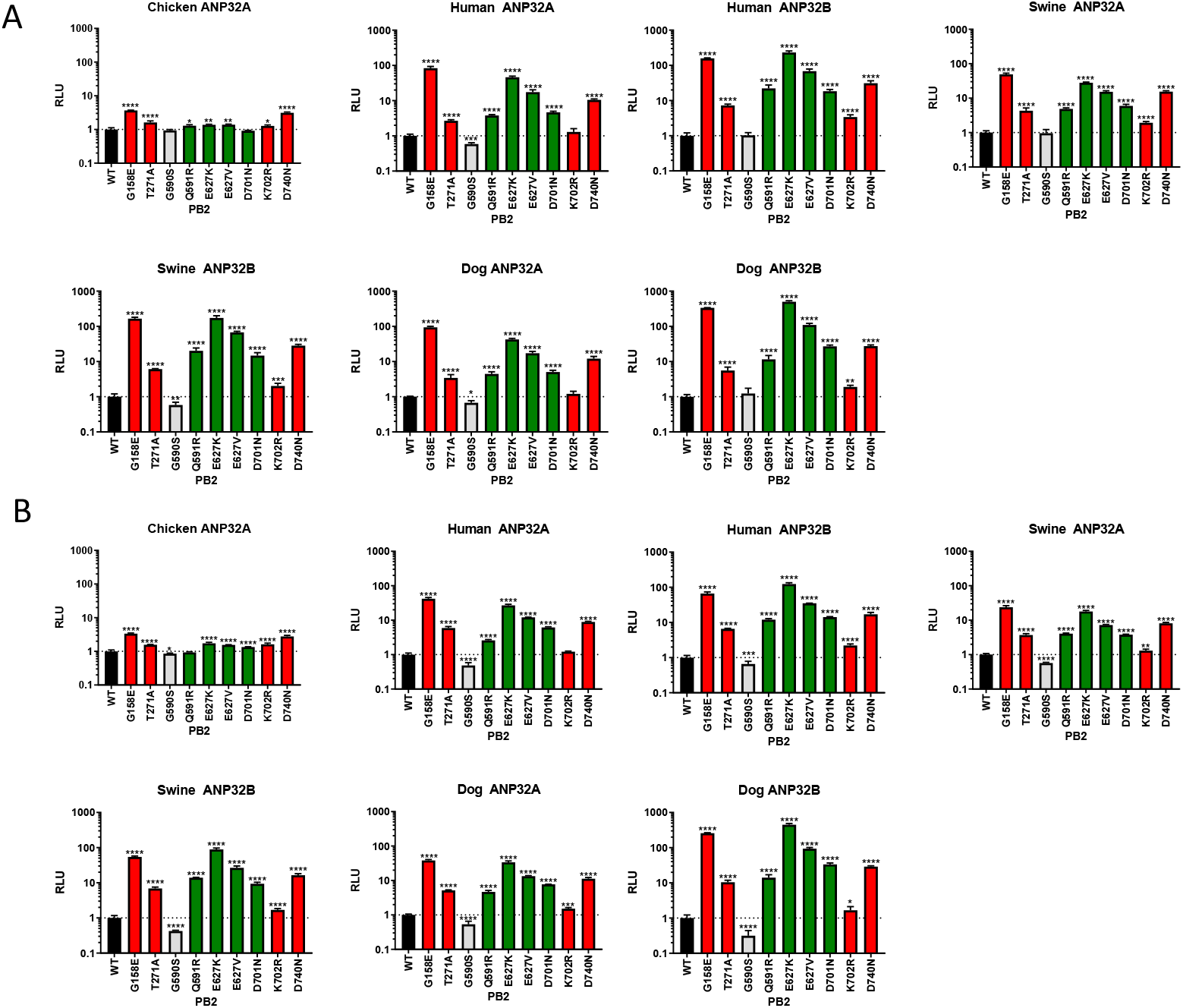
Extended version of Figure 1. Minigenome assays performed in (A) human eHAP dKO cells or (B) chicken DF-1 AKO cells with avian 50-92 polymerase with different mammalian adaptations transfected in along with different avian or mammalian ANP32 proteins, or empty vector. Data throughout indicates triplicate repeats plotted as the mean and standard deviation. Statistical significance was determined by one-way ANOVA with multiple comparisons against WT on log-transformed data. Lognormality of data was confirmed by Shapiro-Wilk test of normality. *, 0.05 ≥ P > 0.01; **, 0.01 ≥ P > 0.001; ***, 0.001 ≥ P > 0.0001; ****, P ≤ 0.0001.

**Figure S2.**
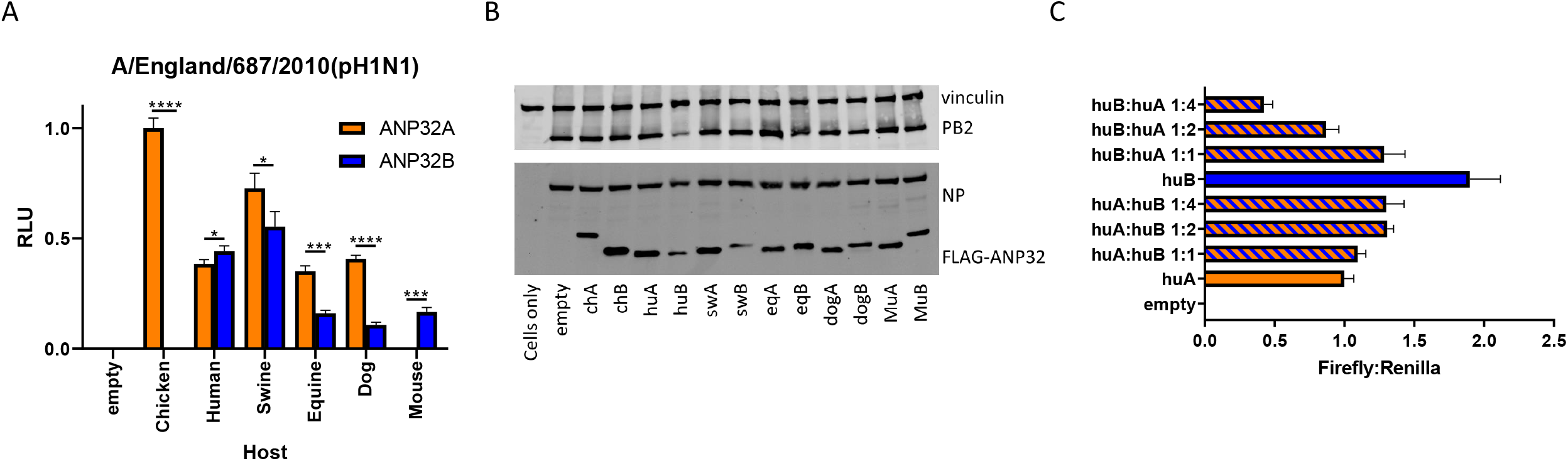
Different mammalian naturally species have ANP32 proteins which are more, or less supportive of influenza virus polymerase. Minigenome assays performed in human eHAP dKO cells with different mammalian polymerases, co-transfected with different mammalian ANP32 proteins expressed (A), different ratios of human ANP32A and B expressed (C). Data throughout indicates triplicate repeats plotted as the mean and standard deviation. Statistical significance was determined by multiple T-tests between different species ANP32A and ANP32B proteins. *, 0.05 ≥ P > 0.01; **, 0.01 ≥ P > 0.001; ***, 0.001 ≥ P > 0.0001; ****, P ≤ 0.0001. (B) Western blot analysis of mutant ANP32 proteins tested in part (A).

**Figure S3.**
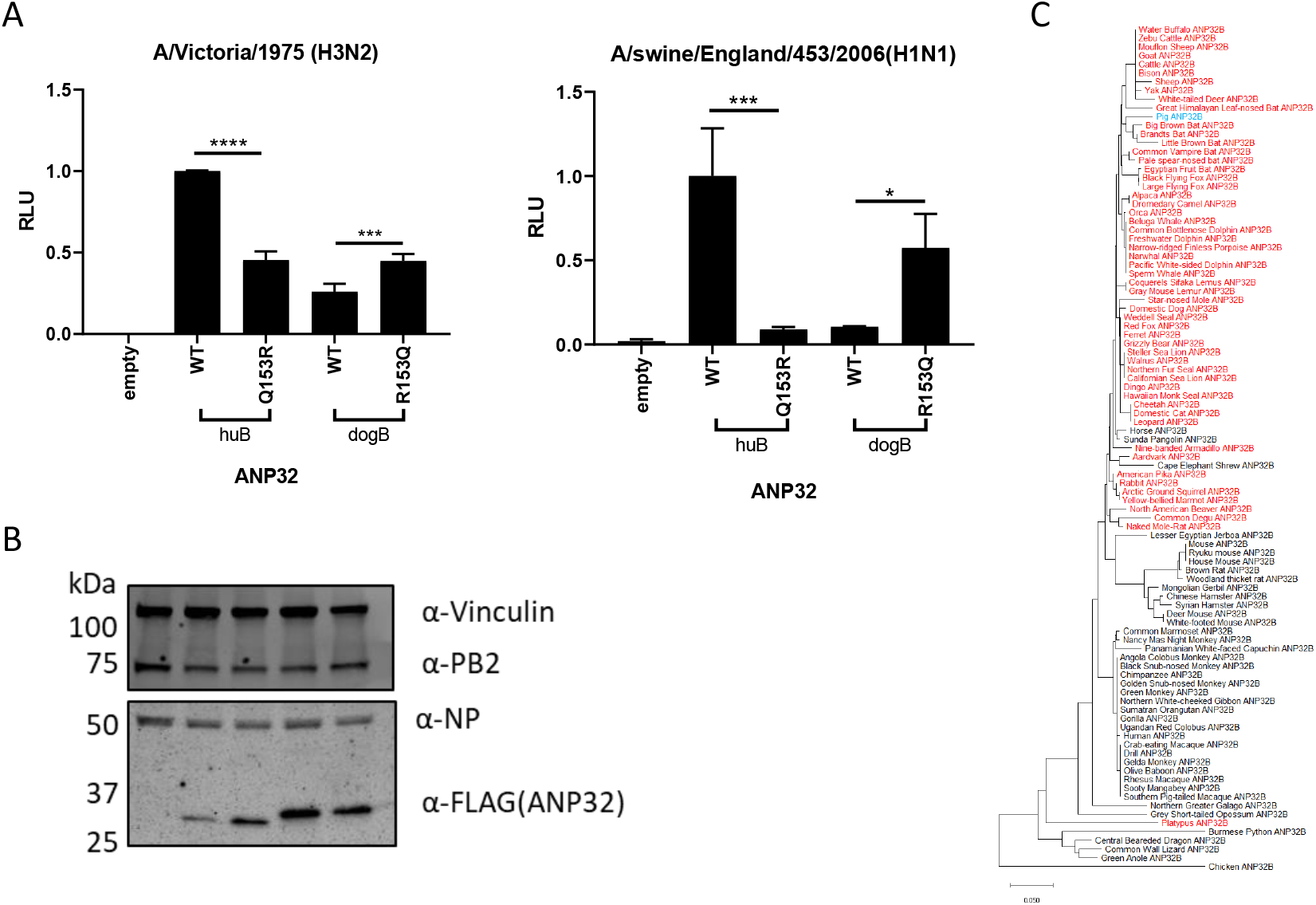
Canine ANP32B poorly supports polymerase activity due to a polymorphism at residue 153. Minigenome assays performed in human eHAP dKO cells with different human or dog ANP32B mutants (A). Data throughout indicates triplicate repeats plotted as the mean and standard deviation. Statistical significance was determined by one-way ANOVA with multiple comparisons, statistical tests performed between WT and mutant ANP32 proteins. *, 0.05 ≥ P > 0.01; **, 0.01 ≥ P > 0.001; ***, 0.001 ≥ P > 0.0001; ****, P ≤ 0.0001. (B) Western blot analysis of mutant ANP32 proteins tested in part (A). (C) Phylogenetic analysis of avian and mammalian ANP32B proteins. Species which contain the weakly proviral signature, 153R shown in red, species with 156Q shown in black, species with 153G shown in cyan. Phylogenetic trees made using the neighbour-joining method based on amino acid sequence.

**Figure S4.**
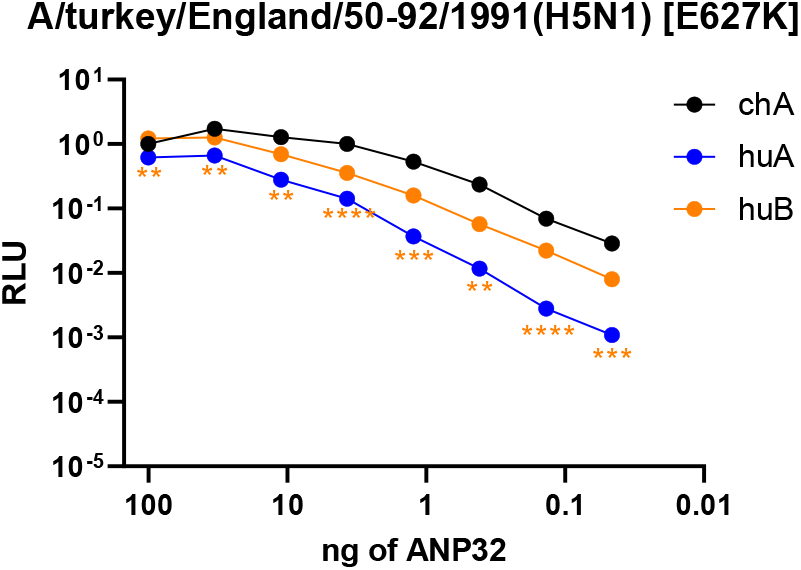
PB2-E627K is more strongly supported by ANP32B across a wide range of plasmid concentrations. Minigenome assays performed in human eHAP dKO cells with chicken ANP32A, or human ANP32A or ANP32B titrated in. Data throughout indicates triplicate repeats plotted as the mean and standard deviation. Statistical significance was determined by multiple T-tests between human ANP32A and ANP32B, statistical tests performed between WT and mutant ANP32 proteins. *, 0.05 ≥ P > 0.01; **, 0.01 ≥ P > 0.001; ***, 0.001 ≥ P > 0.0001; ****, P ≤ 0.0001.

**Figure S5.**
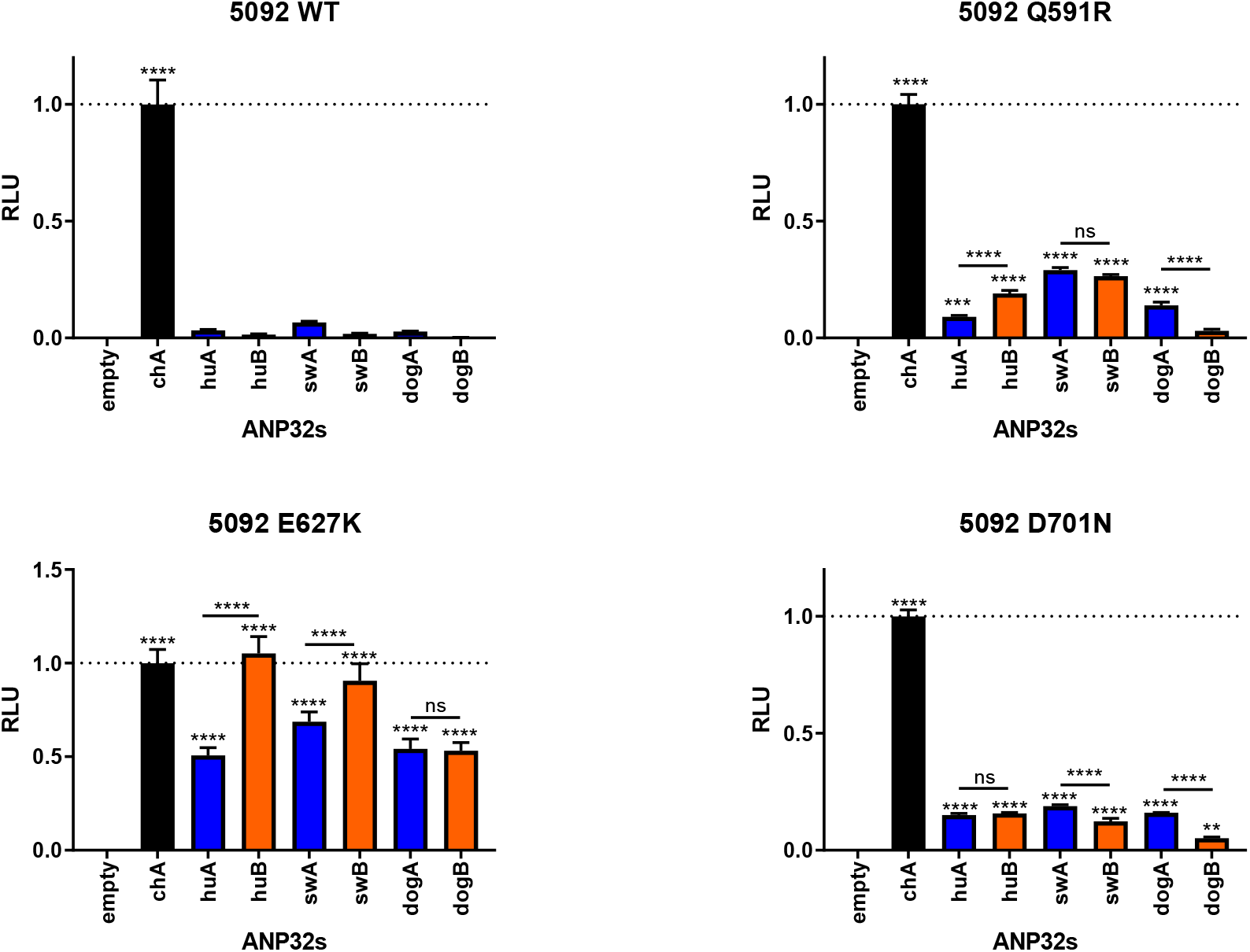
PB2-E627K, but not Q591R or D701N shows a preference for using mammalian ANP32B proteins in DF-1 AKO cells. Data from Figure 1 and Supplementary Figure S1 shown in a different format. Minigenome assays performed in avian DF-1 AKO cells with avian 50-92 polymerase with different mammalian adaptations transfected in along with different avian or mammalian ANP32A (blue bars) or ANP32B proteins (orange bars). Data throughout indicates triplicate repeats plotted as the mean and standard deviation, normalised to chicken ANP32A. Statistical significance was determined by one-way ANOVA with multiple comparisons, statistical tests without comparison bars indicate a comparison against empty vector and between ANP32A and ANP32B proteins from the same species. *, 0.05 ≥ P > 0.01; **, 0.01 ≥ P > 0.001; ***, 0.001 ≥ P > 0.0001; ****, P ≤ 0.0001.

